# Bacteriophage cocktail shows no toxicity and improves the survival of *Galleria mellonella* infected with *Klebsiella* spp.

**DOI:** 10.1101/2023.12.13.571586

**Authors:** Lucy Kelly, Eleanor Jameson

## Abstract

*Klebsiella* spp. are causative agents of healthcare associated infections in patients who are immunocompromised and use medical devices. The antibiotic resistance crisis has led to an increase in infections caused by these bacteria, which can develop into potentially life-threatening illness if not treated swiftly and effectively. Thus, new treatment options for *Klebsiella* are urgently required. Phage therapy can offer an alternative to ineffective antibiotic treatments for antibiotic-resistant bacteria infections. The aim of the present study was to produce a safe and effective phage cocktail treatment against *K. pneumoniae* and *K. oxytoca*, both in liquid *in vitro* culture and an *in vivo Galleria mellonella* infection model. The phage cocktail was significantly more effective at killing *K. pneumoniae* and *K. oxytoca* strains compared with monophage treatments. Preliminary phage cocktail safety was demonstrated through application in the *in vivo G. mellonella* model: where the phage cocktail induced no toxic side effects in *G. mellonella*. In addition, the phage cocktail significantly improved the survival of *G. mellonella* when administered as a prophylactic treatment, compared with controls. In conclusion, our phage cocktail was demonstrated to be safe and effective against *Klebsiella* spp. in the *G. mellonella* infection model. This provides a strong case for future treatment for *Klebsiella* infections, either as an alternative, or adjunct to, antibiotics.

## Introduction

*Klebsiella* spp. are often found as causative agents of healthcare associated infections, mainly causing urinary tract infections (UTIs) and pneumonia [1, 2]. *Klebsiella* have previously been estimated to cause 8% of the total number of nosocomial bacterial infections across Europe and the United States, with *Klebsiella pneumoniae* and *Klebsiella oxytoca* the most prevalent clinically relevant species [3, 4]. Of particular concern are *Klebsiella* infections of immunocompromised patients and those who require the use of medical devices such as catheters and ventilators, because infections in this group of patients have mortality rates as high as 80% [1, 5]. Gram-negative ESKAPE pathogens, such as *K. pneumoniae*, are currently considered to present the greatest bacterial threat to healthcare, as the emergence of antimicrobial resistant strains, resistant to most, or all, available antibiotics is increasing globally [6]. *Klebsiella*, particularly *K. pneumoniae*, accumulate and disseminate multi-drug resistance determinants and are resistant to a wide range of antibiotic classes [7]. Over the past two decades, the prevalence of multi-drug resistant strains of *Klebsiella* has increased throughout the healthcare system, which has led to UTIs, pneumonia and sepsis which are increasingly difficult to treat using the antibiotics currently available [8, 9]. Current approaches used to tackle such infections often focus on the use of a combinatorial approach of antibiotic therapy, however, such treatment has reported a limited efficacy in clinical trials and may be nephrotoxic [10, 11].

Preventing the spread and treating infections caused by antibiotic resistant strains of *Klebsiella* is a challenging prospect, therefore, alternative treatments to traditional antibiotic therapy, such as bacteriophage (phage) therapy, may provide a promising strategy to target these bacteria. A previously published systematic review of multiple phage therapy studies against ESKAPE pathogens reported that phage therapy was safe and effective for treatments caused by these pathogens [12]. The aforementioned study reported that phage therapy was effective at degrading biofilms, reducing bacterial burden, encouraging wound healing and improving patient outcomes. However, despite the urgent requirement for an alternative to antibiotic treatment for *Klebsiella* infections, there are no commercially available phage therapeutics to treat this bacterium.

Phage cocktails are a combination of phage administered as a mixture for phage therapy, as opposed to monophage therapy which uses a single phage [13]. Phage cocktails present a number of advantages compared with monophage therapy, such as an increased host range and decreased resistance rates [13]. Phage cocktails have demonstrated efficacy compared with monophage therapy in an *in vivo* study of *Klebsiella* infection, with the phage cocktail reducing the emergence of phage-resistant mutants and reducing overall bacterial load [14]. A previous study reported the successful use of a phage cocktail to treat a patient with a long-term UTI caused by a multi-drug resistant strain of *K. pneumoniae* [15]. The majority of the available data on the clinical efficacy of phage therapy against *Klebsiella* infections comes from administering phage as a last resort treatment and these compassionate care cases have shown promise, eliminating a chronic UTI, removing *Klebsiella* from the gut and enhancing wound healing [15–18]. A meta-analysis conducted by Al-Anany *et al* on the efficacy of phage therapy for UTIs reported ∼ 72% of studies showed patients improved following phage therapy, while ∼ 99% of patients reported no adverse effects, which suggests that phage therapy is both effective and safe for the treatment of UTIs [19]. However, these studies used personalised phage treatment, there is little currently known about how the efficacy of standardised phage cocktail treatment varies across a wider range of *Klebsiella* strains in *in vivo* models of infection, as most currently published studies focus on a single strain of bacteria.

The aim of the present study was to develop a safe and effective *Klebsiella* phage cocktail against a panel of *K. pneumoniae* and *K. oxytoca* strains and examine the efficacy of this treatment in an *in vivo* invertebrate model of *Klebsiella* infection, using *Galleria mellonella* (waxworm moth) larvae.

## Results

*Virulence index.* The Vi of each individual phage, in addition to the phage cocktail (PhC), was determined against the panel of *Klebsiella* strains, to analyse if the PhC was more effective at killing *Klebsiella* compared with monophage therapy treatment (Fig. 1). The Vi of the PhC against *Kp*30104 was significantly higher compared with phages 7, 10, 12 and 67. Furthermore, the PhC was significantly more effective at killing *Kp*170723 compared all of the monophage treatments tested. The PhC was significantly more effective at killing *Ko*170748 compared with phages 7, 10, 44 and 64. There was no significant difference observed between the Vi of the PhC and single phage against *Kp*13442 and *Kp*171266. None of the phage tested in this study were previously shown to infect *Kp*13442 [20].

**Figure 1.**
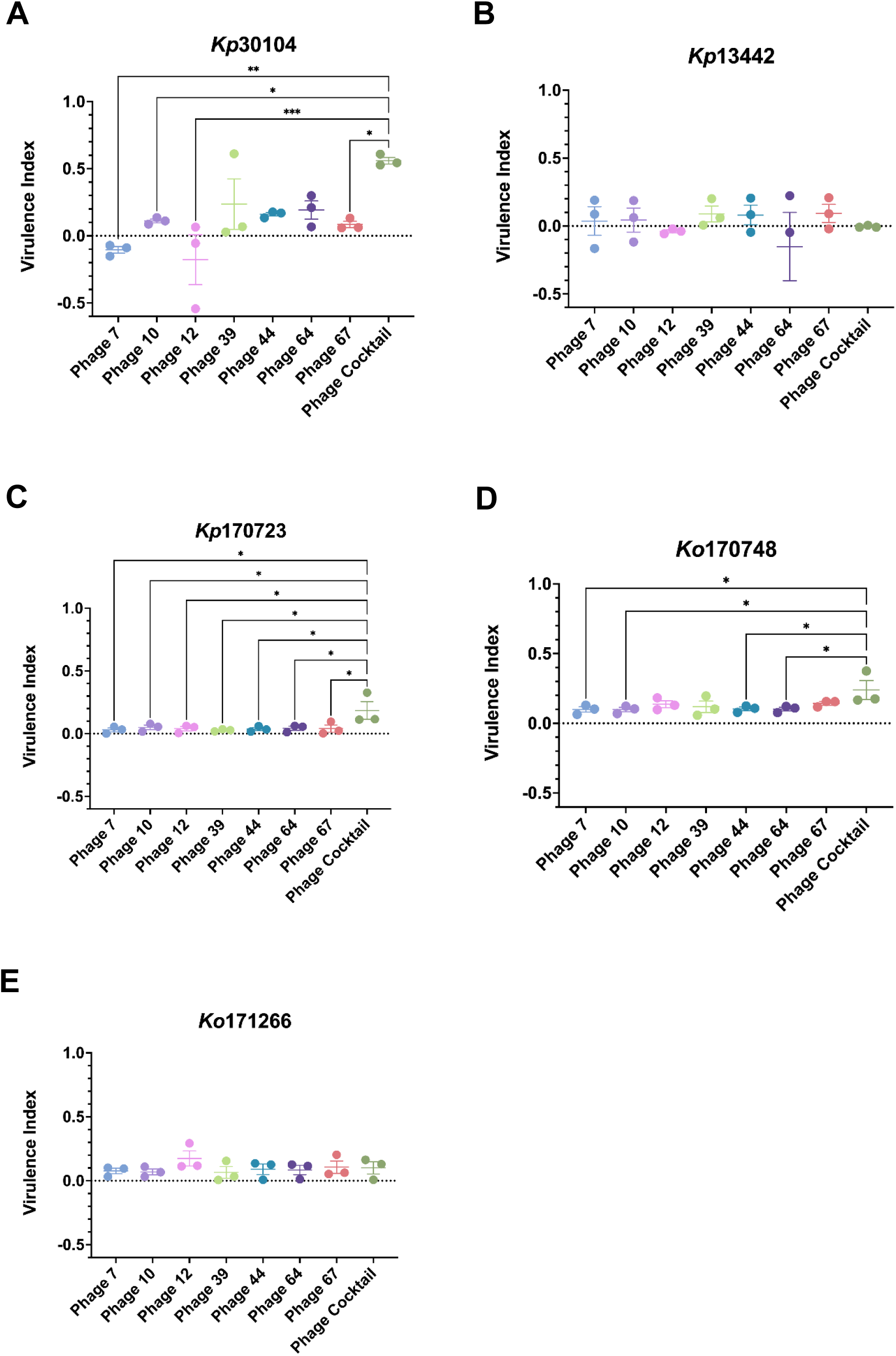
Virulence index of single phage and phage cocktail against. (A) *Kp*30104, (B) *Kp*13442, (C) *Kp*170723, (D) *Ko*170748 and (E) *Ko*171266. *P<0.05, **P<0.01, ***P<0.001 vs. phage cocktail.

*Endotoxin testing and removal*. Endotoxin testing is an important step in the process of PhC preparation as endotoxin present in phage preparations is a highly immunogenic compound which can cause endotoxic shock when present at high quantities [21]. The concentration of endotoxin present in the PhC before and after endotoxin removal is presented in Table 1. The maximum accepted level of endotoxin present in a medicinal product is 0.2 EU/kg/h for intrathecal administration or 5 EU/kg/h for intravenous administration, which is equivalent to 1.6×10^3^ EU/10^9^ PFU of phage treatment [22–24]. As the level of endotoxin present in the PhC preparation after endotoxin removal was 70.41 EU/10^9^ PFU, the PhC preparation was deemed safe for use in the *G. mellonella* model.

**Table 1.**
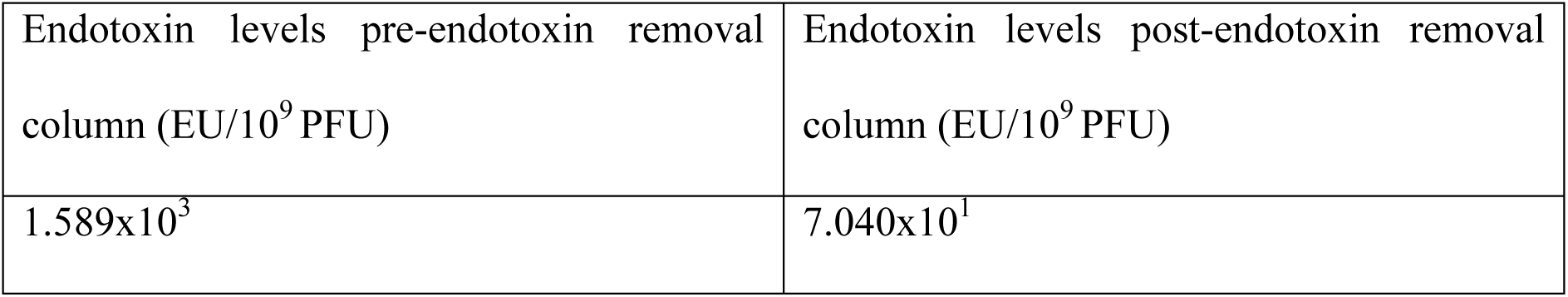
Concentration of endotoxin in phage cocktail preparation before and after endotoxin removal.

| Endotoxin levels pre-endotoxin removal<br>column (EU/10<sup>9</sup> PFU) | Endotoxin levels post-endotoxin removal<br>column (EU/10<sup>9</sup> PFU) |
| --- | --- |
| 1.589x10<sup>3</sup> | 7.040x10<sup>1</sup> |

*LD_50_ of Klebsiella in G. mellonella larvae*. The survival of *G. mellonella* larvae injected with different *Klebsiella* strains was monitored for 5 days to determine the LD_50_ dose of each bacterial strain (Fig. 2). Based on these survival curves, the LD_50_ of each of the *K. pneumoniae* and *K. oxytoca* strains was calculated (Table 2). This showed the optimal dose of *K. pneumoniae* 2×10^4^-5×10^5^ and *K. oxytoca* was 1×10^6^ to obtain LD_50._

**Figure 2.**
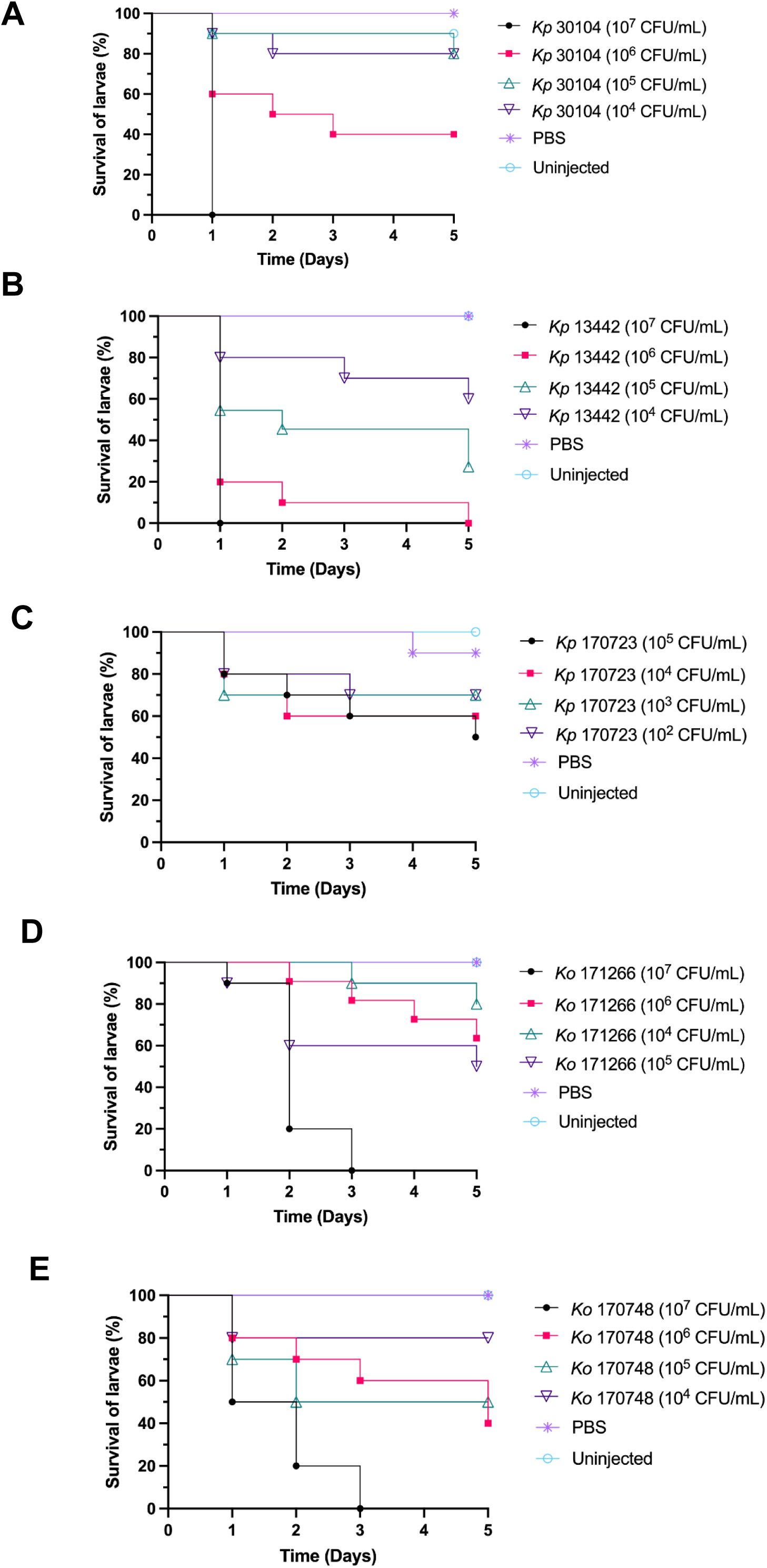
Survival curves of LD_50_ doses of. (A) *Kp*30104, (B) *Kp*13442, (C) *Kp*170723, (D) *Ko*170748 and (E) *Ko*171266 in *Galleria mellonella* larvae after 5 days.

**Table 2.** LD_50_ dose of different *Klebsiella* strains in *Galleria mellonella*.

| Strain | LD<sub>50</sub> (CFU/ml) |
| --- | --- |
| *Kp30104* | 5x10<sup>5</sup> |
| *Kp13442* | 2x10<sup>4</sup> |
| *Kp170723* | 2x10<sup>5</sup> |
| Ko170748 | 1x10<sup>6</sup> |
| Ko171266 | 1x10<sup>6</sup> |

*Survival of* G. mellonella *infected with* Klebsiella *with prophylactic phage therapy.* Prophylactic PhC phage therapy was administered to larvae to determine if this treatment regime was effective at rescuing *G. mellonella* larvae from infection and death caused by *Klebsiella* infection. Prophylactic PhC was administered to the larvae 4 h before infection with *Klebsiella* and survival of the larvae was monitored for 5 days (Fig. 3). The survival of larvae prophylactically treated with PhC (MOI=1) and infected with *Kp*30104 was significantly increased, compared with bacteria-only controls. Prophylactic PhC (MOI=1, 10 and 100) of *Kp*170723– and *Ko*170748*-*infected larvae significantly increased survival compared with bacteria-only controls. The survival of larvae prophylactically treated with PhC (MOI=10 and 100) and infected with *Ko*171266 was significantly increased, compared with bacteria-only controls. There was no significant difference observed in survival of phage-injected larvae compared with controls, which indicated that the PhC did not induce death in these larvae.

**Figure 3.**
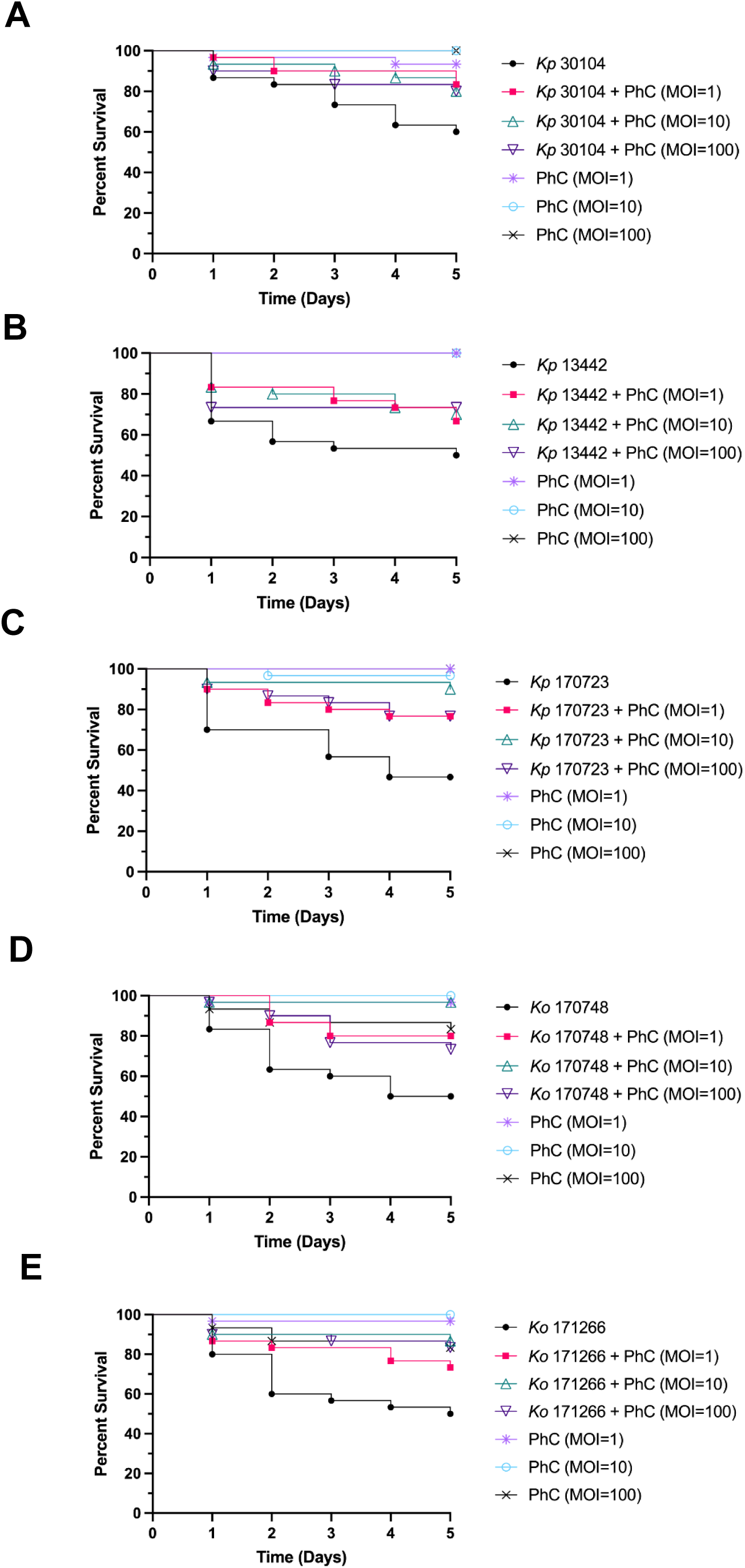
Survival curves of prophylactic phage cocktail treatment against *Galleria mellonella* infected with. (A) *Kp*30104, (B) *Kp*13442, (C) *Kp*170723, (D) *Ko*170748 and (E) *Ko*171266.

*Survival of G. mellonella infected with Klebsiella treated with co-injection phage therapy*. Co-injection of *Klebsiella* and PhC was administered to larvae to determine if this treatment regime could prevent infection and death of larvae injected with *Klebsiella*. PhC and *Klebsiella* were combined and injected into larvae and survival of *G. mellonella* was monitored for 5 days (Fig. 4). Co-injection of PhC (MOI=1, 10 and 100) significantly improved survival of larvae injected with *Kp*30104, compared with bacteria-only controls. The survival of larvae treated with PhC (MOI=100) and *Ko*170748 was significantly improved compared with bacteria-only controls. There was no significant difference observed in survival of phage-injected larvae compared with controls, which indicated that the PhC did not induce death in these larvae.

**Figure 4.**
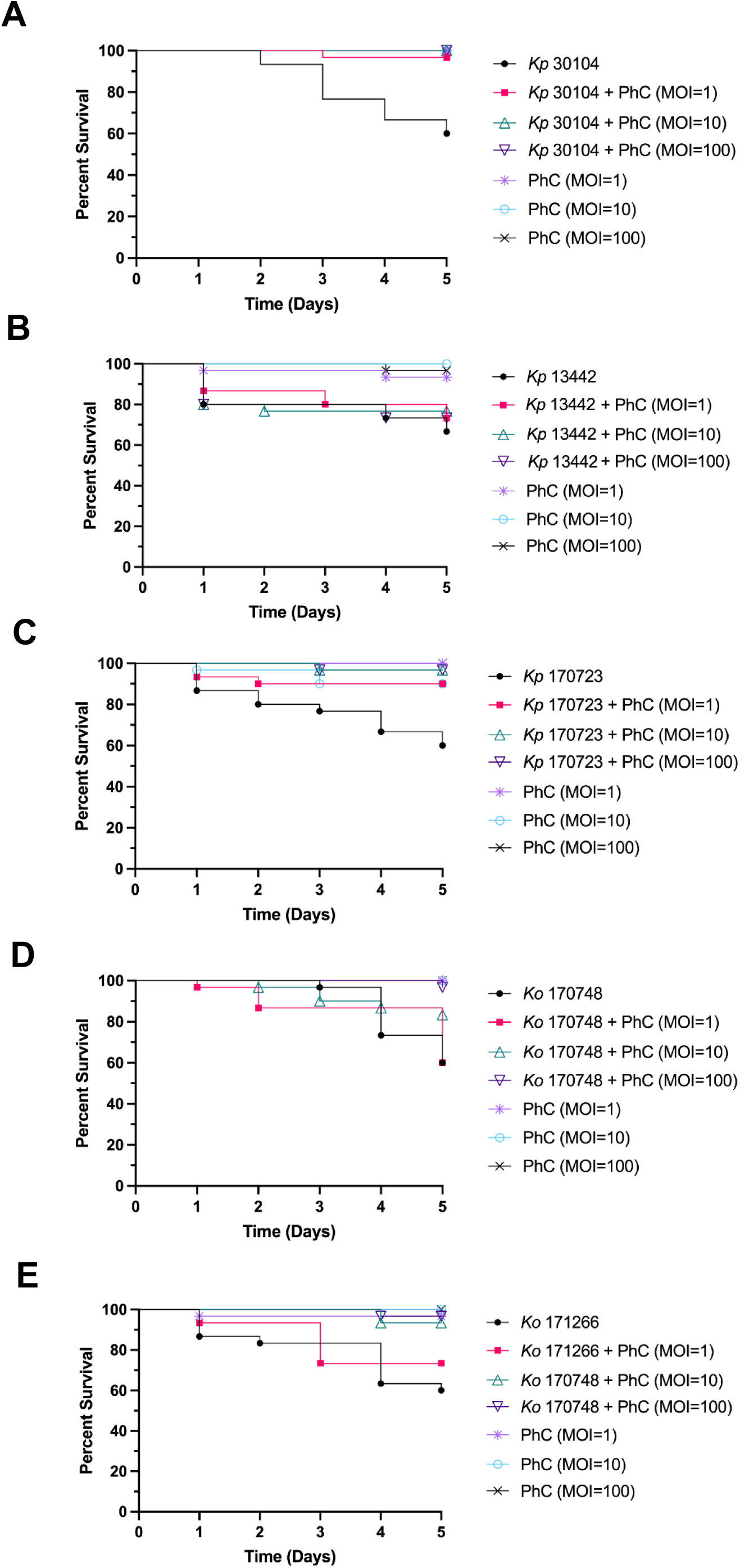
Survival curves of co-injection phage cocktail treatment against *Galleria mellonella* infected with. (A) *Kp*30104, (B) *Kp*13442, (C) *Kp*170723, (D) *Ko*170748 and (E) *Ko*171266.

*Survival of G. mellonella infected with* Klebsiella *treated with remedial phage therapy.* Remedial phage therapy with the PhC was administered to *G. mellonella* larvae in order to analyse if this treatment regime was effective at preventing death caused by *Klebsiella* infection. The PhC was administered 4 h following *Klebsiella* injection of the larvae and survival of *G. mellonella* was monitored for 5 days (Fig. 5). Remedial PhC (MOI=10) significantly improved the survival of larvae injected with *Kp*30104, compared with bacteria-only controls. There was no significant difference observed in survival of phage-injected larvae compared with controls, which indicated that the PhC did not induce death in these larvae.

**Figure 5.**
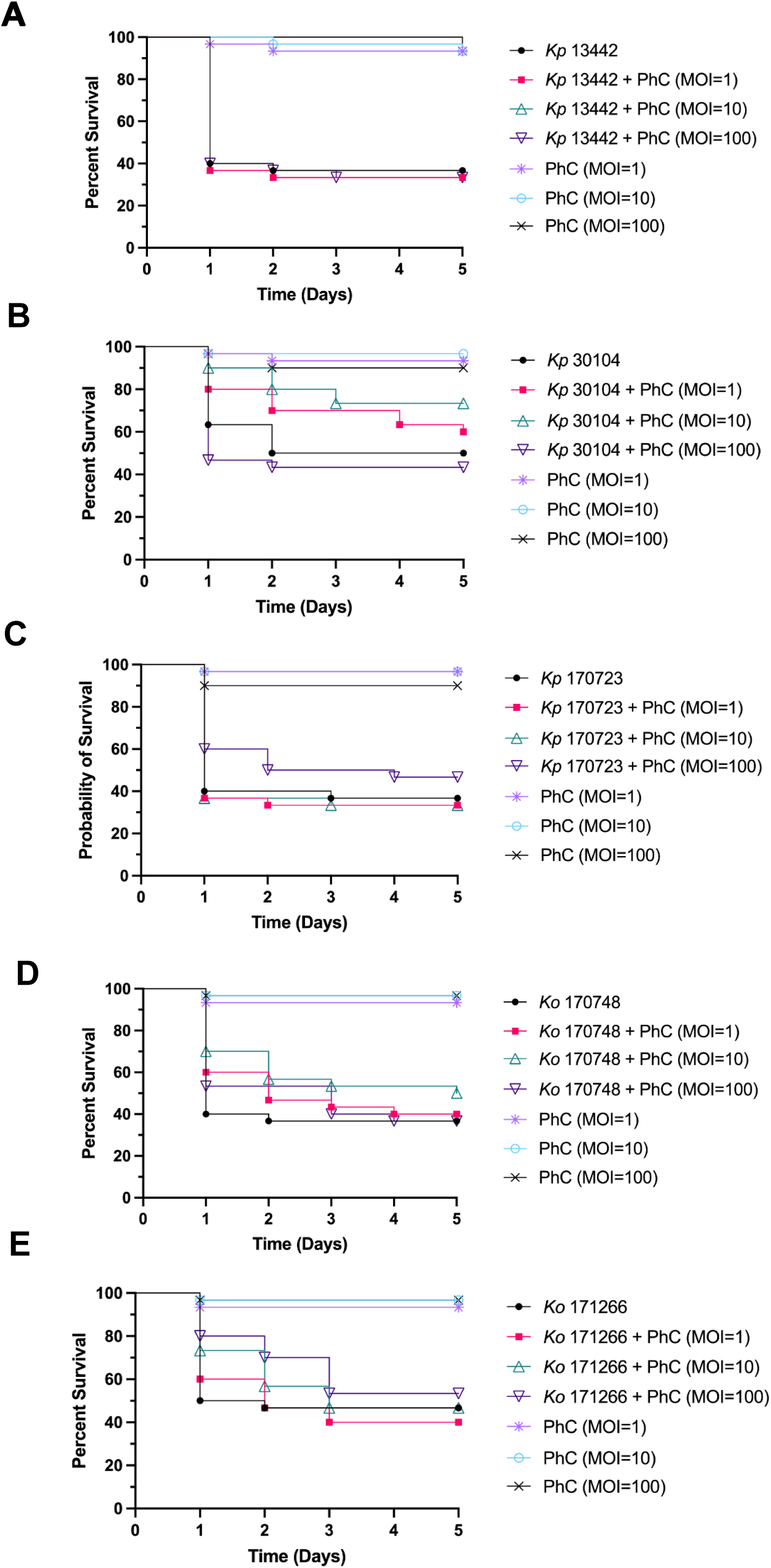
Survival curves of remedial phage cocktail treatment against *Galleria mellonella* infected with. (A) *Kp*30104, (B) *Kp*13442, (C) *Kp*170723, (D) *Ko*170748 and (E) *Ko*171266.

## Discussion

Our study confirms the efficacy of a phage cocktail (PhC) treatment against *K. pneumoniae* and *K. oxytoca* strains in a *G. mellonella* infection model with prophylactic phage therapy, showing no adverse effects. A number of previously published studies have characterised effective monophage or PhC treatments against a single strain, sequence type or capsule type of *Klebsiella*, but few have produced an effective and safe PhC treatment against multiple strains and species of *Klebsiella* [25–29]. To the best of our knowledge, no studies have previously reported on the efficacy of a PhC treatment on both culture collection and clinically-isolated strains of *Klebsiella*.

The phage selected for inclusion into the PhC treatment were genetically distinct and had the widest host range available from a panel of previously characterised phages [20]. This PhC consisted of phages that spanned six phylogenetic groups, five families, three subfamilies, six genera, four sources of isolation and six capsule type targets [20]. This strategy was selected for design of the PhC, as a combination of genetically distinct phages are more likely to be effective at removing target bacteria compared with monophage treatment, if a PhC contains phage with different host receptor targets, the host is unlikely to gain multiple mechanisms of resistance simultaneously without significantly impairing its own fitness [13]. Our PhC was effective at killing four out of five *Klebsiella* strains tested, with the exception of *Kp*13442, a strain resistant to all individual phages tested [20]. *Kp*13442 was taken tested in this study to determine if a PhC treatment would demonstrate synergy against a phage-resistant strain of bacteria under *in vivo* conditions, however, this was not the case. The Vi results demonstrated that the PhC was more effective at killing *Kp*30104, *Kp*170723 and *Ko*170748 than a number of the monophage treatments tested, which suggested that the PhC has the potential to be a more effective treatment against these *Klebsiella* strains than monophage therapy. The PhC was the most effective against *Kp*170723 compared with the single phage treatments, which indicated synergistic killing effects between the phages in the cocktail, despite no monophage treatment demonstrating a high Vi against *Kp*170723.

Of the three PhC phage therapy dosing regimes administered here, the most effective at killing a range of *Klebsiella* strains in the invertebrate model was prophylaxis, followed by co-injection. Prophylactic phage therapy was effective at increasing the survival of *G. mellonella* infected with *Kp*170723 and *Ko*170748, two clinically-isolated strains of *Klebsiella*, which highlighted the potential for this PhC to be used to treat clinical infections caused by these strains. This phage therapy regime was also effective at increasing survival of *Kp*30104. Prophylactic phage therapy could potentially act as a type of intervention therapy, preventing disease in patients when used prior to surgery or in patients who require the use of medical devices. Prophylactic phage therapy aims to prevent colonisation and infection by bacteria before they can cause harm, as opposed to treating an established infection [30–32]. This treatment regime presents an attractive option to tackle healthcare associated *Klebsiella* infections, as these are often a result of the contamination of medical devices, such as catheters and ventilators [33]. Co-injection of phage and bacteria is indicative of a clinical situation whereby phage therapy is administered to a patient at the same time the patient may potentially be colonised by *Klebsiella*, for example upon admission to hospital or the use of a medical device. Co-injection of the higher doses of PhC in the present study were effective at increasing the survival of *G. mellonella* injected with *Kp*30104 and *Kp*170723. The success of this co-injection regime of phage therapy may potentially be due to an increased frequency of bacteria-phage interactions, as the phage were combined directly with bacteria before injection into the larvae. This may enable the phage to more rapidly adhere to the invading bacteria, negating the need for chance encounters between phage and bacteria during systemic circulation in the larvae, to enable phage infection, proliferation and killing of the target bacteria [34]. Remedial phage therapy is a clinically-relevant scenario for PhC administeration to a patient with an ongoing *Klebsiella* infection. In the present study, remedial phage therapy was the least successful regime tested, however, one concentration of the PhC was effective at increasing the survival of larvae infected with *Kp*30104. Therefore, remedial PhC phage therapy shows potential, but requires further optimisation of the dose, timing or PhC formulation to maximise efficacy. This observed lower efficacy of remedial phage therapy compared with prophylactic or co-injection phage therapy is consistent with previously published studies on the efficacy of different phage therapy regimes in *G. mellonella* [35, 36].

A previous study on the use of a prophylactic PhC against several bacterial species, including *K. pneumoniae*, was effective against infections in a mouse model [37]. The pretreatment of medical devices has previously been reported to be highly effective at reducing colonisation of devices by pathogens and prevent biofilm formation [38]. The data presented in the present study adds to a growing body of literature that supports the efficacy of prophylactic phage therapy for the prevention of bacterial infections. Furthermore, this demonstrates that phage can persist in an invertebrate host, were not degraded by the *G. mellonella* innate immune system or inactivated through binding to organic matter. Therefore, prophylactic phage therapy, at the treatment concentration provided in the present study, circulated throughout the *G. mellonella* system and encountered host bacteria.

The higher doses of PhC therapy administered were most effective at increasing the survival of *G. mellonella* compared with the lower doses of PhC. This association has also been reported in previous studies of phage therapy in *G. mellonella* [35, 39, 40]. This is potentially due to increased bacteria-phage interactions that may occur as a result of an increased concentration of phage compared with low-dose phage therapy [13].

One of the aims of the present study was to design a safe PhC for use in an *in vivo* model of *Klebsiella* infection. The survival of the control phage-only groups of larvae were comparable to the untreated and injection trauma controls in each phage therapy regime, which demonstrated that the PhC was not innately toxic and could be carried forward safely into more complex infection models, such as mammalian models of *Klebsiella* infection. The acquisition of preliminary safety data is an important part of phage therapy design, as patient safety is an essential pillar for the introduction of phage therapy in human clinical trials [41].

Further development of this PhC treatment could include the testing of multiple doses of PhC therapy, as a single dose of the PhC was tested in the present study, as multiple doses of PhC may potentially increase the efficacy of the treatment. Additionally, the impact of PhC as an adjunct to antibiotic treatment is an important consideration for future phage therapy studies. Phage-antibiotic synergy has previously been reported and has shown promise in treating bacterial infections, including those caused by *K. pneumoniae* [42–44].

To conclude, the present study demonstrates the design of a safe and effective PhC treatment against *K. pneumoniae* and *K. oxytoca*. The PhC was effective at increasing the survival of *G. mellonella* infected with both culture collection and clinically-isolated strains of *Klebsiella*. Therefore, the PhC presented in this study shows potential for the future treatment of *Klebsiella* infections, as an alternative or adjunct to antibiotic treatments.

## Materials and methods

*Bacterial culture*. The details of the bacterial strains used in this study are presented in Table 3. *Klebsiella* strains were maintained on Lysogeny broth (LB; Merck) agar plates. As required, liquid cultures were prepared by inoculating 10 ml of LB with a single colony of bacteria and incubating overnight at 37°C and 180 rpm shaking.

**Table 3.** Details of *Klebsiella* strains used in the present study.

| Strain name | Origin | Isolation source | Capsule type (KL) |
| --- | --- | --- | --- |
| *Klebsiella pneumoniae* 30104<br>(Kp30104) | DSMZ culture collection | Human blood | KL3 |
| *Klebsiella pneumoniae* 13442<br>(Kp13442) | NCTC culture collection | Hospital | KL110 |
| *Klebsiella pneumoniae* 170723<br>(Kp1707233) | Clinical isolate | Urine | KL2 |
| *Klebsiella oxytoca* 170748 (Ko170748) | Clinical isolate | Catheter specimen urine | O1v1 |
| *Klebsiella oxytoca* 171266 (Ko171266) | Clinical isolate | Urostomy urine | OL104 |

*Bacteriophage propagation*. The details of the bacteriophage used in this study are presented in Tables 4 and 5 and the host range of these phage is presented in Table 6. A 10 µl volume of phage lysate was added to a log-phase culture of the *Klebsiella* host strain in LB supplemented with 5 mM CaCl_2_ (Thermo Fisher Scientific) and 5 mM MgCl_2_ (Thermo Fisher Scientific) and incubated at 37°C with 180 rpm shaking overnight. Phage lysate was centrifuged at 3,220 x g for 20 min at room temperature and the supernatant filtered through a 0.2 µm syringe filter (Sarstedt) to remove host cells. Phage were enumerated through serial dilution, then mixing 50 µl of the diluent with 125 µl of log-phase host bacteria, followed by incubation at room temperature for 10 min. Each dilution was mixed with 625 µl of LB top agar (0.4% agar) supplemented with 5 mM CaCl_2_ and 5 mM MgCl_2_ and immediately plated on 6-well plates containing solidified 1% LB agar. The solidified overlay plates were incubated at 37°C overnight and phage plaques enumerated.

**Table 4.** Details of bacteriophage strains used in the present study.

| Phage name | Lab ID | Isolation source | Isolation strain | Capsule target (KL) | Accession no. |
| --- | --- | --- | --- | --- | --- |
| *Klebsiella* | 7 | Slurry, | *Klebsiella* | - | PRJEB40160 |
| phage vB<br>KaS-Ahsoka |  | slurry tank,<br>UK | *aerogenes* DSM<br>30053 |  |  |
| *Klebsiella*<br>phage vB<br>KppS-Totoro | 10 | Estuary,<br>Jelitkowo,<br>Poland | *Klebsiella*<br>*pneumoniae*<br>DSM 30104 | KL3 | PRJEB40166 |
| *Klebsiella*<br>phage vB<br>KoM-Pickle | 12 | Estuary,<br>Jelitkowo,<br>Poland | *Klebsiella*<br>*oxytoca* DSM<br>25736 | KL74 | PRJEB40176 |
| *Klebsiella*<br>phage vB<br>KqM-Weterburg | 39 | Raw<br>sewage,<br>Spernal<br>sewage<br>works, UK | *Klebsiella*<br>*quasipneumoniae*<br>DSM 28211 | KL35 | PRJEB40173 |
| *Klebsiella*<br>phage vB<br>KqP-Goliath | 44 | Raw<br>sewage,<br>Spernal<br>sewage<br>works, UK | *Klebsiella*<br>*quasipneumoniae*<br>DSM 700603 | KL53 | PRJEB40163 |
| *Klebsiella*<br>phage vB<br>KpM-Wobble | 64 | Mixed<br>liquor<br>sewage,<br>Spernal<br>sewage<br>works, UK | *Klebsiella*<br>*pneumoniae*<br>170958 | KL28 | PRJEB40182 |
| *Klebsiella* | 67 | Mixed | *Klebsiella* | KL14 | PRJEB40178 |
| *phage* vB<br>KpM-KalD |  | liquor<br>sewage,<br>Spernal<br>sewage<br>works, UK | *pneumoniae*<br>DSM 13439 |  |  |

**Table 5.**
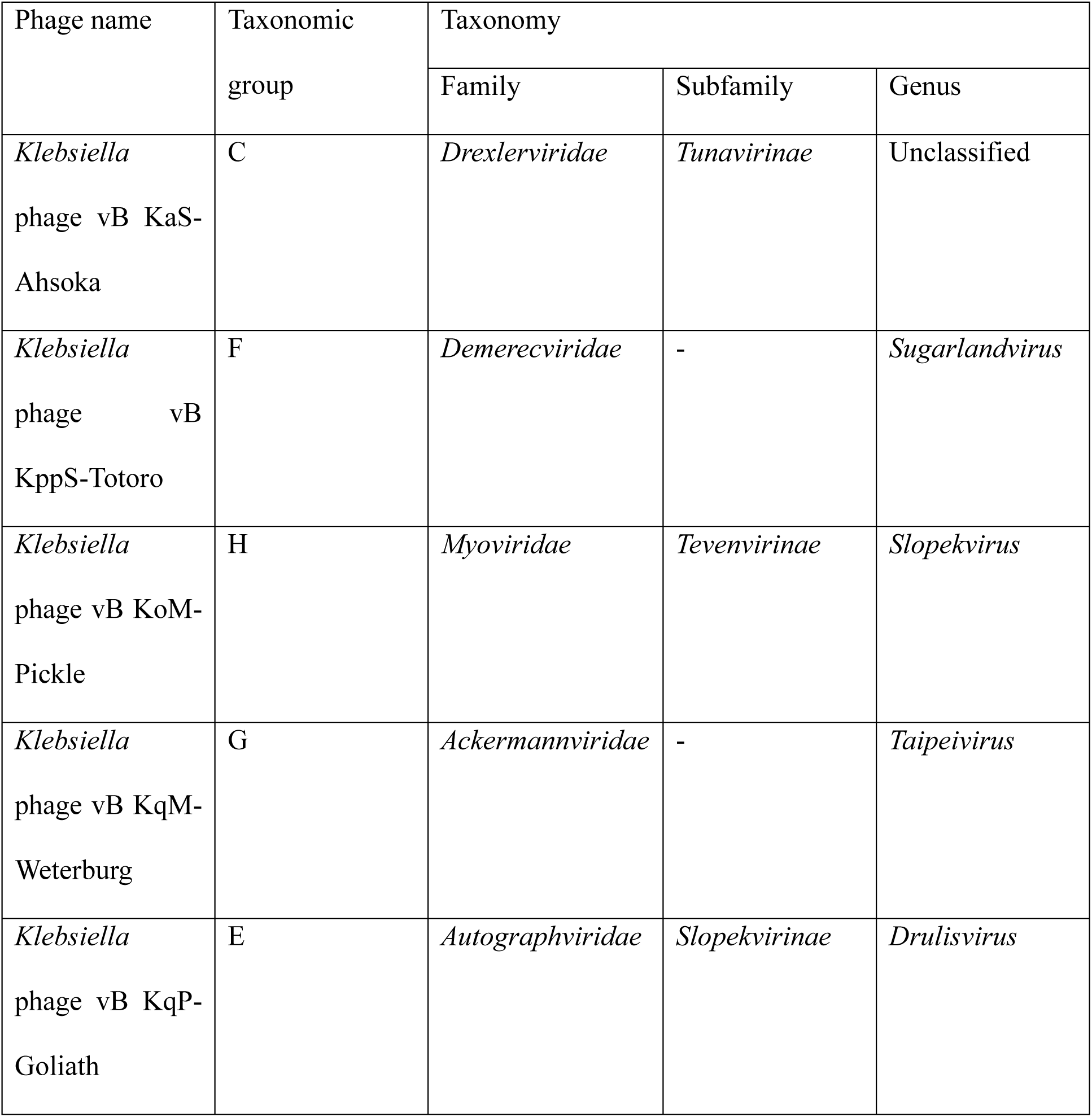
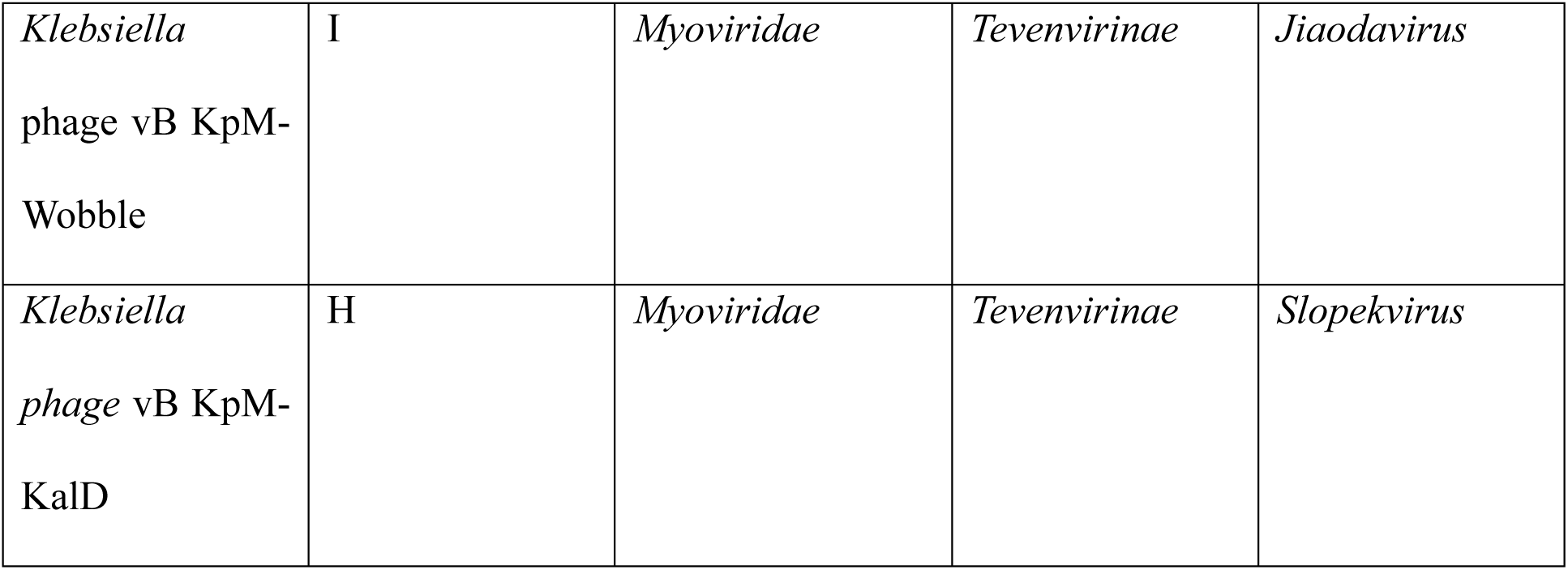
Taxonomy of phage used in the present study.

**Table 6.** Host range of phage used in the present study.

| *Klebsiella*<br>strain | Phage 7 | Phage 10 | Phage 12 | Phage 39 | Phage 44 | Phage 64 | Phage 67 |
| --- | --- | --- | --- | --- | --- | --- | --- |
| *Kp30104* | Yes | Yes | Yes | Yes | - | Yes | Yes |
| *Kp13442* | - | - | - | - | - | - | - |
| *Kp170723* | - | - | Yes | Yes | Yes | Yes | Yes |
| *Ko170748* | - | Yes | Yes | - | - | - | Yes |
| *Ko171266* | - | Yes | Yes | - | - | - | Yes |

*Bacteria-phage growth curves.* The Virulence Index (Vi) was used to quantify the efficacy of single phage and the phage cocktail against the *Klebsiella* strains in this study [45]. An overnight culture of *Klebsiella* was refreshed 1:100 in LB and incubated at 37°C and 180 rpm shaking until log-phase growth was achieved. Phage were serially diluted from an MOI of 1 to 10^-7^ and 10 µl of each phage dilution was added to 190 µl of *Klebsiella* in a 96-well plate. The optical density of the samples was measured at 600 nm every 10 min for 18 h at 37°C using a FLUOstar Omega Microplate Reader (BMG Labtech). The Vi of each sample was then calculated.

*Endotoxin testing and removal*. Endotoxin testing of phage lysates was performed using the Pierce™ Chromogenic Endotoxin Quant Kit (Thermo Fisher Scientific, Inc.) was used according to the manufacturer’s instructions. The Pierce™ High Capacity Endotoxin Removal Spin Columns (Thermo Fisher Scientific, Inc.) were used to remove endotoxin from phage lysates.

*Maintenance of* G. mellonella. *G. mellonella* larvae (LiveFood UK Ltd.) were obtained, stored at 4°C immediately upon arrival and used within 2 days of delivery. Larvae were individually weighed and those with a weight between 0.20-0.30 g were selected for further experiments. Groups of 10 larvae were randomly assigned to each treatment condition and stored in Petri dishes throughout the experiment.

*Determining the LD_50_ of* Klebsiella *strains in* G. mellonella. The LD_50_ of each *Klebsiella* strain was calculated over a 5-day period. An overnight culture of *Klebsiella* was diluted 1:100 in LB and incubated at 37°C and 180 rpm until log-phase growth was achieved. Cultures were centrifuged twice at 3,220 x g for 15 min at room temperature, cell pellets washed with PBS and the final pellet re-suspended and serially diluted in PBS. *G. mellonella* injection sites were sterilized using 70% ethanol. A 10 µl dose of *Klebsiella* was injected into the last left proleg of the larvae using a 30 G needle on a Hamilton 500 µl gastight syringe fitted with a Hamilton PB600-1 repeating dispenser. The Kp170723 doses used were 10^5^, 10^4^, 10^3^ and 10^2^ CFU/larvae. The doses for the remaining four *Klebsiella* strains were 10^7^, 10^6^, 10^5^ and 10^4^ CFU/larvae. Negative controls of PBS-injected and untreated larvae were included in each experiment. Larvae were incubated at 37°C in the dark for 5 days and remained unfed. Larval survival was monitored daily, and larvae were scored as either live or dead every 24 h, with larvae classified as dead once they turned black and immobile. Dead larvae were removed each day. At the end of the experiments, larvae were euthanized by placing them in a –20°C freezer for <2 h, then moved to a –80°C freezer for overnight storage before disposal. The LD_50_ dose calculated was used to infect larvae in subsequent experiments.

*Phage therapy of Klebsiella-infected G. mellonella*. Three different PhC phage therapy regimens (prophylaxis, co-injection and remedial treatment), along with bacteria– and phage-only controls and PBS-injected and untreated larvae were prepared. A 10 µl dose of *Klebsiella* was used in each of the bacteria-treated larvae. PhC dilutions were prepared in PBS to an MOI of 1, 10 or 100 for each *Klebsiella* strain to be tested. A 10 µl dose of PhC was used in each of the phage-treated larvae.

The phage therapy regimens were designed as follows: i) Prophylaxis, PhC injected at 0 h and *Klebsiella* injected at 4 h; ii) co-injection, PhC and *Klebsiella* were mixed and immediately injected at 0 h; and iii) remedial, *Klebsiella* injected at 0 h and PhC injected at 4 h. In the treatment groups where two injections were required (prophylaxis and remedial therapy), the first injection was in the last left proleg and the second injection was in the last right proleg. The survival of larvae was monitored over 5 days.

*Statistical analysis*. Statistical analysis and data visualisation was performed using GraphPad Prism (version 9.5.1; Dotmatics). One-way analysis of variance with Tukey’s post-hoc test was used to determine statistically significant differences between multiple groups. Kaplan-Meier survival curves were produced and a Log-rank Mantel-Cox test performed to determine statistical significance between survival curves. Data are presented as the mean ± standard deviation of three independent experiments. P<0.05 was considered to indicate a statistically significant difference.

## Acknowledgements

The funders had no role in study design, data collection and interpretation, or the decision to submit the work for publication. This work was partially supported by the MIBTP DTP PhD studentship awarded to LK.

